# In vivo visualization of nitrate dynamics using a genetically encoded biosensor

**DOI:** 10.1101/2022.08.29.505680

**Authors:** Yen-Ning Chen, Heather Cartwright, Cheng-Hsun Ho

## Abstract

Nitrate (NO_3_^-^) uptake and distribution are critical to plant life. Although the upstream regulation of nitrate uptake and downstream responses to nitrate in a variety of cells have been well-studied, it is still not possible to directly visualize the spatial and temporal distribution of nitrate with high resolution at the cellular level. Here, we report a nuclear-localized, genetically encoded biosensor, nlsNitraMeter3.0, for the quantitative visualization of nitrate distribution in *Arabidopsis thaliana*. The biosensor tracked the spatiotemporal distribution of nitrate along the primary root axis and disruptions by genetic mutation of transport (low nitrate uptake) and assimilation (high nitrate accumulation). The developed biosensor effectively monitors nitrate concentrations at cellular level in real time and spatiotemporal changes during the plant life cycle.

**One-Sentence Summary:** A genetically encoded biosensor for in vivo visualization of spatiotemporal nitrate levels at a cellular resolution.

## Main Text

The plant root is essential to nutrient uptake. Nitrate (NO_3_^-^), is a major nitrogen source and is one of the most limiting factors in agricultural production (*1, 2*). Within the root, nitrate levels differ dramatically between root cell types (*3, 4*). Under nitrate limitation, plants can optimize morphological and physiological parameters, for example, root growth can be directed towards nutrient deposits in the soil, the root surface area can be locally increased, or the transporter density on the membrane can be altered. Moreover, metabolic conversion, storage and translocation of nitrogen compounds are modified (*5, 6*). In order to adjust these parameters, plants have to monitor both the external and intracellular nitrate concentrations to determine NO_3_^-^ acquisition needs by plant roots.

NO_3_^-^ uptake predominantly occurs from the soil/rhizosphere into roots. Once in a root cell, NO_3_^-^ ions can diffuse within the symplasm from cell to cell. NO_3_^-^ ions can serve as an osmotic compound or be assimilated in the root to produce organic nitrogen for cellular growth either locally or be loaded into xylem vessels for transport to the shoot (*7*)., NO_3_^-^ uptake, the rate of NO_3_^-^ acquisition by the plant, depends on the surface area of the root, and the environmental factors that affect root growth will also affect NO_3_^-^ capacity. Furthermore, the root system is very plastic, and NO_3_^-^ availability itself strongly affects root development. However, we still do not fully understand the most fundamental aspects about NO_3_^-^ uptake by plant roots, such as: which tissue(s) is(are) responsible for NO_3_^-^ uptake? is NO_3_^-^ uptake distributed all along the root? and, is NO_3_^-^ uptake restricted to specific zones only? In addition, the exact intercellular path from the outer root layers towards the central stele is only hypothesized and not experimentally proven. It has proven difficult to track nitrate molecules within the plant tissue. Some studies have reported nitrate detection, however, most of these techniques either lack spatial resolution, e.g. radioactive isotope (*8, 9*) and the Griess method (*10*), or have limitations to their use, e.g. vibrating electrodes (*11, 12*), positron-emitting tracer imaging (*13, 14*), or secondary ion mass spectrometry (*15*).

Other ions have been monitored in living tissue through Förster Resonance Energy Transfer (FRET)-based biosensors. FRET sensors are fusion proteins that report on a target molecule through interactions with a sensory domain that cause changes in a protein conformation (*16*). These conformational changes affect the efficiency of energy transfer from a fused FRET donor fluorescent protein to a fused FRET acceptor fluorescent protein. Changes in energy transfer can be detected by measuring changes in the relative intensity of the two fluorescent proteins (ratio change) after excitation of the donor; the ratio change reports target molecule concentration. We report here the development of a biosensor to monitor the dynamics of NO_3_^-^ in plants.

The bacterial NasR protein is a soluble receptor that contains the NIT (nitrate- and nitrite-sensing) domain, which serves as an NO_3_^-^ binding pocket (*17–19*). We generated a biosensor based on a FRET biosensor by cloning the NIT domain as a Gateway Entry clone and then recombining it with a previously designed Gateway destination vector (pDR-FLIP39) that carries enhanced dimerization (ed) variant of Aphrodite (edAFP), as the FRET acceptor, and of enhanced cyan fluorescent protein (edeCFP) as the FRET donor (*20*). The fusion proteins were expressed in protease-deficient yeast, purified (*20*), and analyzed in a spectrofluorometer for NO_3_^-^-dependent alterations in the fluorescence emission curves after FRET donor excitation at 428 nm (donor excitation (Dx); Fig. S1). Within the NIT domain fusion protein, the fluorophores were within Förster distance, as evidenced by resonance energy transfer; however, NO_3_^-^ addition did not trigger a significant change in the energy transfer rate between the emission at 530 nm (donor excitation acceptor emission, DxAm) and the emission at 488 nm (donor excitation donor emission, DxDm) that could act as a FRET ratio change sensor (ΔDxAm/DxDm). The initial emission ratio (ΔDxAm/DxDm) of the NIT domain fusion protein was greater than 1.2 (Fig. S1). To further optimize the sensor, we tested sensor replacing the NIT domain with the entire NasR protein (Fig. 1A). The NasR fusion construct showed a NO_3_^-^-triggered increase in energy transfer between donor emission at 530 nm (DxAm) and acceptor emission at 488 nm (DxDm). The NasR FRET biosensor was named <u>Ni</u>tra<u>Met</u>er<u>1.0</u> (NiMet1.0) and reports nitrate levels with a positive ratio change (ΔDxAm/DxDm) (Fig. S2). Fluorescent protein pair variants and different lengths of linkers can have dramatic effects on sensor responses (*21–23*). In attempts to optimize NiMet1.0, different FRET pairs including brightness variants and truncation variants and different lengths of linkers to either the N- or C-terminus of the Gateway Destination vector [pDR-FLIP30, pDR-FLIP39, and pDR-FLIP42-linker (*20*)] were tested. NiMet2.0 (Fig. S3), a FRET pair variant containing Citrine and mCerulean, was consistently NO_3_^-^ responsive. NasR with L12 linkers showed a larger NO_3_^-^-triggered response when fused to the Citrine/mCerulean pair (Fig. S3). Furthermore, RasR with no L12 linkers sandwiched by Aphrodite.t9 and mCerulean (pDR-FLIP30) yielded the highest ratio change and the lowest FRET initiation ratio (NiMet3.0; Fig. 1B). With the information of the crystal structure of NasR (*17*) and our observed DxAm/DxDm values for NiMet3.0 (hereafter referred to as NiMet3.0 emission ratio) with and without NO_3_^-^ (Fig. 1C), one hypothesis is that NiMet3.0 switches from a low FRET to high FRET average state upon binding to NO_3_^-^.

**Fig. 1.**
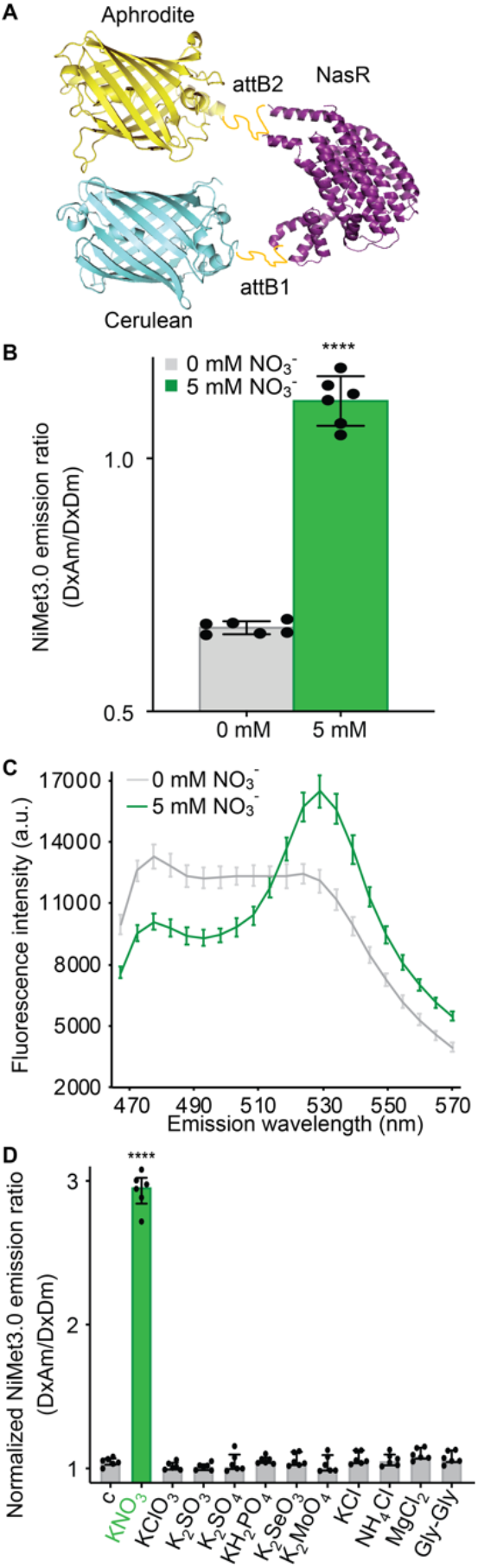
Engineering and specificity for NO_3_^-^ of nitrate biosensor-NiMet 3.0. **A**, Structural model of NiMet 3.0 bound to NO_3_^-^. NasR, an NO_3_^-^ binding protein, was fused via *attB1* and *attB2* linkers to a fluorescent protein FRET pair (donor, Aphrodite, and acceptor, Cerulean). The NasR protein (purple) representation is from a published structure of NasR [PDB 4AKK(*17*)]. The Aphrodite (yellow) representation is from a published structure of Venus [PDB 1MYW(*35*)] and the Cerulean (blue) representation is from a published structure of Cerulean [PDB 2WSO(*36*)]. **B and C**, Fluorescence emission ratio at 530 nm (B) and emission wavelength scan (C) of purified NiMet3.0 protein with and without NO_3_^-^. Nitrate concentration as indicated in figures. **D**, Substrate specificity of purified NiMet3.0 treated with the indicated compounds at 5 mM concentrations. Only nitrate triggered responses that were significantly different from control, c (*, *p* <0.0001, *t*-test). The presented data are Mean ± s.d. of six biological repeats. Experiment performed as in Fig. 1B.

To test the specificity of NiMet3.0 to NO_3_^-^, different forms of nitrogen and other anions were examined. Neither other anions nor other nitrogen forms, like ammonium or a peptide, triggered emission ratio changes, thus the NiMet3.0 sensor is specific to NO_3_^-^ (Fig. 1D). To determine the dynamic range of NO_3_^-^ detection by NiMet3.0, we measured the dissociation constant (*K*_d_) of purified NiMet3.0 *in vitro* by tracking dosedependent changes in NiMet3.0 emission ratios for NO_3_^-^ (Fig. 2A). The sensitivity of NasR for NO_3_^-^ is in the micromolar to millimolar range (*19*). The *K*_d_ of NiMet3.0 was ~90 μM for NO_3_^-^ and reach a maximum at NO_3_^-^ concentrations above 1 mM (Fig. 2B). This affinity is comparable with the NasR sensitivity for NO_3_^-^. Non-responsive variants of NiMet3.0, an important control of NiMet3.0 specificity, were generated via mutation of NasR residues involved in NO_3_^-^ binding (Fig. 2B). NiMet3.0-R49A, -R50A, -R176A, and -R236A carry alanine substitutions in the predicted NO_3_^-^ binding pocket of NasR based on the crystal structure of the NasR protein and that have been shown to disrupt NO_3_^-^ responses (*17*). NiMet3.0-R49A and -R236A still showed detectable response to NO_3_^-^, but with lower emission ratios compared with NiMet3.0, whereas NiMet3.0-R50A and -R176A, the substitution in the NasR binding pocket, showed no responses to NO_3_^-^, likely due to disrupted salt-bridges that function in the interaction with NO_3_^-^ (Fig. 2C) (*17*). The above mutant biosensors are evidently non-responsive to NO_3_^-^ and, carry a nitrate binding pocket that is predicted to be non-responsive *in planta* with endogenous NO_3_^-^. Together, these data strongly support the hypothesis that NiMet3.0 specifically measures NO_3_^-^ concentrations and can report dynamics of changes in NO_3_^-^ levels.

**Fig. 2.**
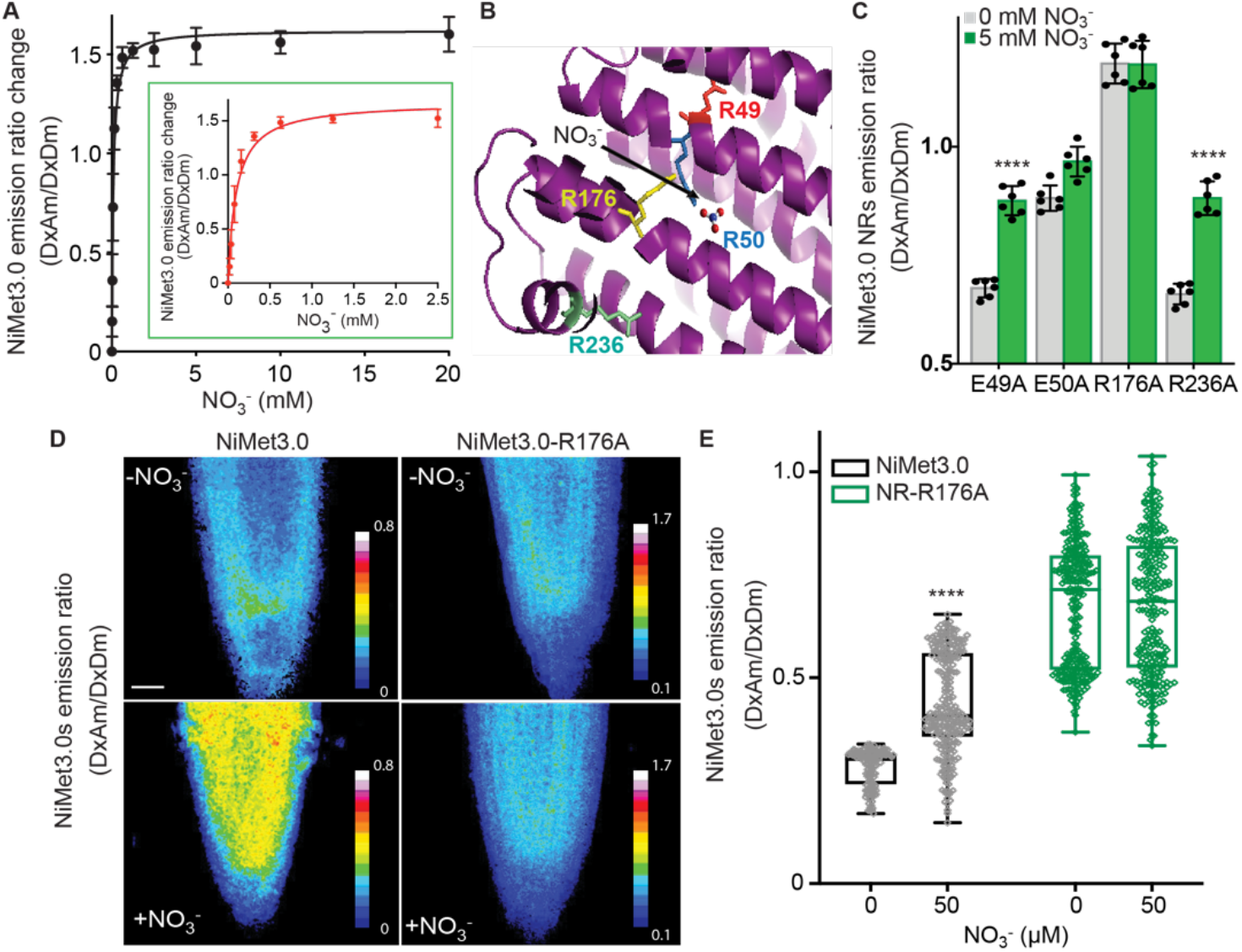
Fluorescence emission ratio response of purified NiMet3.0 to NO_3_^-^ *in vitro* and *in vivo*. **A**, NiMet3.0 fluorescence response to increasing concentrations of NO_3_^-^. Inset: Enlargment of the NiMet3.0 fluorescence response from 0 to 2.5 mM NO_3_^-^. **B**, NiMet3.0 residues of the nitrate binding pocket of NasR mutagenized to make NitMet3.0 non-responsive constructs (NiMet3.0 NR). Four residues of NasR, R49, R50, R176, and R236 (red, blue, yellow and green, respectively) were mutagenized to Ala. **C**, Fluorescence emission ratios of purified NiMet3.0-NR proteins with and without NO_3_^-^ treatment. Nitrate concentration as indicated in figures. Student’s *t*-test \*\*\*\**P* value < 0.0001. Mean and s.d. of six biological repeats are presented. **D,** Images of NiMet3.0 and NiMet3.0 NR-R176A emission ratios in 6-day-old roots in transgenic Col-0 grown with or without 5 mM NO_3_^-^. **E**, Corresponding quantitative analysis of NiMet3.0s emission ratios of root in D. Beeswarm box plot of NiMet3.0s emission ratios from root region (*n* > 80 areas from three independent seedlings for each genotype of three biological experiments). Student’s *t*-test \*\*\*\**P* value < 0.0001. NiMet3.0 emission ratios were statistically different compared to no nitrate.

To test specificity of the NiMet3.0 response to NO_3_^-^ *in planta*, stable transgenic *Arabidopsis* lines expressing either NiMet3.0 or the non-responsive control NiMet3.0-R176A (under the control of the strong constitutive CaMV35S) were generated. Transgenic lines expressing NiMet3.0, but not NiMet3.0-R176A, showed significant emission ratio changes to NO_3_^-^ in roots (Fig. 2D; quantification in Fig. 2E), indicating that NiMet3.0 can specifically detect NO_3_^-^ in plants.

A sensor targeted to the nucleus allows the analysis of NO_3_^-^ accumulation in the compartment most relevant to endogenous NO_3_^-^ perception and facilitates the comparison of nlsNiMet3.0 emission ratios between nearby cells. To assess the control of NO_3_^-^ distribution *in planta*, we generated stable transgenic *Arabidopsis* lines expressing a nuclear-targeted variant of NiMet3.0 (nlsNiMet3.0) under the control of a promoter fragment previously shown to direct broad expression [p16 (*24*)]. Purified nlsNiMet3.0 showed similar *in vitro* responses to NO_3_^-^ as NiMet3.0 (Fig. S4). Expression of nlsNiMet3.0 did not result in detectable phenotypic changes in seedlings or plants (Fig. S5). Through recording the emission ratios of primary root cells exposed to NO_3_^-^ pulses, nlsNiMet3.0 showed a rapid response to NO_3_^-^ pulses, and its signal was reversible when NO_3_^-^ withheld (Fig. 3A). In roots, the *K*_d_ of nlsNiMet3.0 was ~130 μM for NO_3_^-^ (Fig. 3B and Fig. S6). This affinity is comparable with the NiMet3.0 affinity for NO_3_^-^ *in vitro*.

**Fig. 3.**
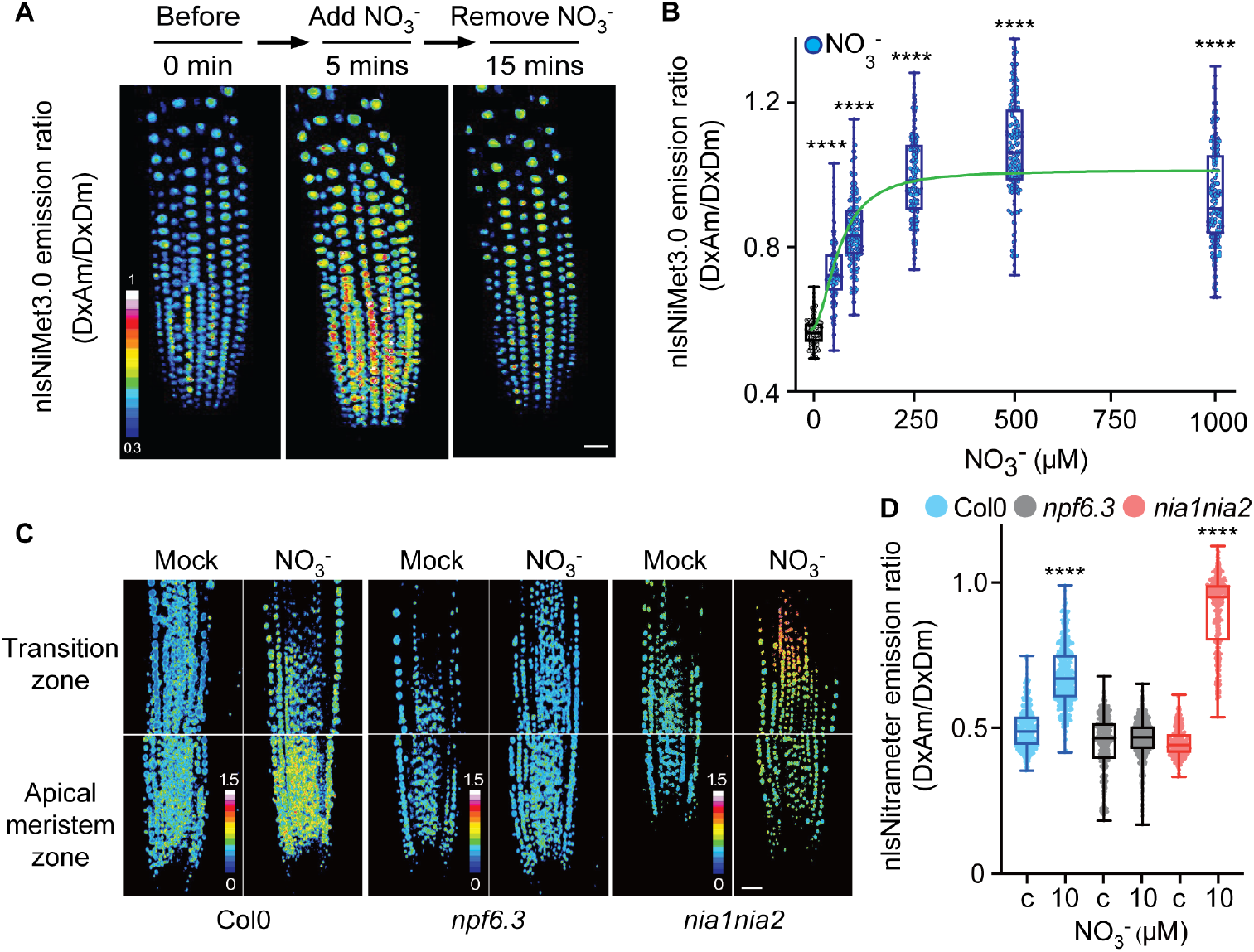
Emission ratios of nlsNiMet3.0 in root tips before and after NO_3_^-^ treatment in Arabidopsis roots. **A**, Three-dimensional images of nlsNiMet3.0 emission ratios of 5-day-old roots in transgenic Col-0 before an NO_3_^-^ pulse, after the NO_3_^-^ pulse, and after removing the NO_3_^-^. 50 μM of NO_3_^-^ was used. **B**, Beeswarm and box plot of NO_3_^-^ concentration-dependent nlsNiMet3.0 emission ratios for nuclei of root tips from Fig. S6. Green line indicates as nonlin fit of nlsNiMet3.0 *K*_d_ curve. Student’s *t*-test \*\*\*\**P* value < 0.0001. Mean and s.d. of three biological repeats are presented. **C**, Images of nlsNiMet3.0 emission ratios of 6-day-old root zones (meristem and transition zone) in FRET transgenics in wild-type Col-0, *npf6.3*, and *nia1nia2* backgrounds grown with or without NO_3_^-^. **D**, Corresponding quantitative analysis of nlsNiMet3.0 emission ratios of root in C. Beeswarm and box plot of nlsNiMet3.0 emission ratios for nuclei of central region (*n* > 180 nuclei from three independent seedlings for each genotype of three biological experiments). nlsNiMet3.0 emission ratios were statistically different in *npf6.3* and *nia1nia2* compared to Col-0 backgrounds (c, as mock as control). **** *P* value < 0.0001.

To explore if NiMet3.0 is suitable for measuring NO_3_^-^ distribution in plants, we investigated nlsNiMet3.0 emission ratios in roots of wild type Col-0, a nitrate transporter mutant [*npf6.3/NTR1;1/chl1-5*, (*25*)], and a nitrate reductase mutant [*nia1nia2*, (*26*)] seedlings were grown on agar plates and exposed to long days (16 h light/8 h dark) for five days (Fig. 3C). NO_3_^-^ uptake into root cells requires nitrate transporters (*25*). In wild-type roots, we observed an overall higher emission ratio in seedlings grown on NO_3_^-^-containing agar compared with those grown without NO_3_^-^. There was an apparent gradient of NO_3_^-^ in the root tip, with high nlsNiMet3.0 emission ratios in the apical meristem zone that reduced to lower nlsNiMet3.0 emission ratios in the root transition zone (Fig. 3C), although local variation was observed. As expected, the nitrate transporter *npf6.3* mutant plants showed lower nlsNiMet3.0 emission ratios in all root zones with or without NO_3_^-^ in the media compared to wild type, supporting the idea that NPF6.3 functions as a major nitrate transporter bringing external NO_3_^-^ into roots. Furthermore, there was an overall increase of nlsNiMet3.0 emission ratios with NO_3_^-^ treatment in the root of the *nia1nia2* mutant compared to wild type and, interestingly, higher nlsNiMet3.0 emission ratios in the cortical cells of the transition zone (Fig. 3C), suggesting an area of higher NIA1/NIA2 protein or activity levels in root. Our findings support the idea that nlsNiMet3.0 is potentially suitable for measuring NO_3_^-^ distribution *in planta*. Corresponding quantification results of Fig. 3C are shown in Fig. 3D.

To explore which tissue(s) or zone(s) along the root are responsible for NO_3_^-^ uptake, we analyzed nlsNiMet3.0 emission ratios in roots of *Arabidopsis* seedlings before and 30 minutes after treatment with exogenous NO_3_^-^. Similar to endogenous NO_3_^-^ treatment shown in Fig. 3C, overall higher nlsNiMet3.0 emission ratios were observed in root meristem and transition zones with exogenously applied NO_3_^-^ (Fig. 4A), although the differential nlsNiMet3.0 emission ratios across the root could result from differential depletion activity. Treatment of Arabidopsis roots did not increase nlsNiMet3.0 emission ratios in endodermis cells, whereas exogenous NO_3_^-^ triggered increased nlsNiMet3.0 emission ratios in epidermis, pericycle, and stele cells, as well as the highest ratios in cortex cells (Fig. 4B). After washing out the exogenous NO_3_^-^, the increased nlsNiMet3.0 emission ratio was rapidly reduced in all root cells. Only the cortex cells in the root meristem maintained relatively high levels after wash-out (Fig. 4A, B). It should be noted that after accumulation of exogenous NO_3_^-^, nlsNiMet3.0 was able to report the depletion of NO_3_^-^ from all types of cells of the roots (Fig. 3A and 4A-B).

**Fig. 4.**
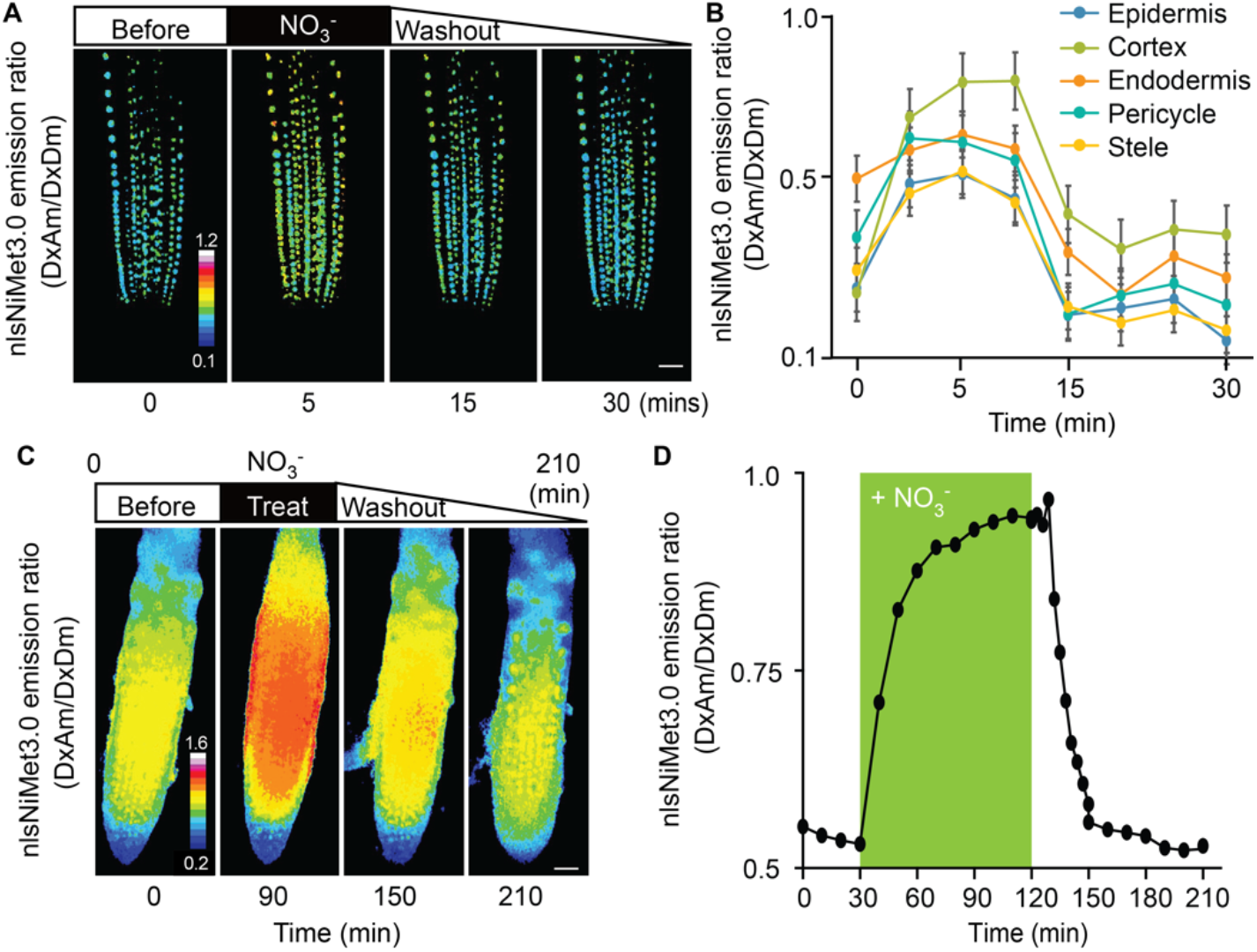
Emission ratios of nlsNiMet3.0 in root under time-course treatment of NO_3_^-^. **A** and **B**, Images and corresponding quantitative analysis of Arabidopsis root before and after NO_3_^-^ treatment. Images obtained after incubation with 50 μM NO_3_^-^ for 5 min. Images were taken before or immediately after NO_3_^-^ treatment for 5 mins, or during the washout at 15 and 30 mins. Scale bar, 25 μm. **B**, Quantitative analysis of nlsNimet3.0 emission ratios for nuclei of epidermis, endodermis, cortex, pericycle, and stele cells in roots from A. Complete experiments were repeated at least three times with similar results. **C**, Time-course treatment of nlsNiMet3.0 with NO_3_^-^. Images were taken before or immediately after NO_3_^-^ treatment for 90 min or at 150 min and 210 min, during the washout. Scale bar, 25 μm. **D,** Corresponding quantitative analysis of nlsNimet3.0 emission ratios of root in C. Complete experiments were repeated at least three times with similar results.

The accumulation of exogenously applied NO_3_^-^, detected by nlsNiMet3.0 in the nuclei of root cells, reflects a balance between import and depletion activities, for example metabolism, export, and compartmentation. To quantify this cooperative activity with high spatio-temporal resolution, we performed time-course experiments on Arabidopsis roots using light-sheet microcopy, a microfluidic device that allows imaging of roots growing in Fluorinated Ethylene Propylene (FEP) tubes with a perfusion control system (Fig. S7). After 90 minutes of perfusion with 10 μM of NO_3_^-^, the nlsNiMet3.0 emission ratio reached its V_max_, indicating that the concentration of NO_3_^-^ accumulation in roots was around 250 μM, reflective of the *K*_d_ of nlsNiMet3.0 we obtained in Arabidopsis root (Fig. 3B). With washout, the emission ratio rapidly reduced back to initial levels (Fig. 4C-D and Movie S1). These results also indicated that the import and depletion activities were dynamically balanced in root after 90 min of perfusion with 10 μM of NO_3_^-^.

Here we report the engineering and analysis of the first biosensor for nitrate (NO_3_^-^) in *planta* and its use to examine the NO_3_^-^ distribution in whole root tissues (Fig. S8). The biosensor nlsNiMet3.0 can be used in living plant roots to quantify NO_3_^-^ concentrations and dynamics.

## Acknowledgments

We gratefully acknowledge Dr. Wolf B. Frommer for discussions and suggestions. We thank Addgene for distributing plasmids donated by Wolf B. Frommer. We thank the Advanced Optics Microscope Core Facility at Academia Sinica for technical support for fluorescence imaging. The core facility is funded by Academia Sinica Core Facility and Innovative Instrument Project (AS-CFII-108-116). We thank Anita K. Snyder for English editing.

## Author contributions

Conceptualization: C.-H.H.

Methodology: C.-H.H., Y.-N.C., H.N.C.

Investigation: C.-H.H., Y.-N.C., H.N.C.

Writing, C.-H.H.

Supervision, C.-H.H.

## Data and materials availability

All data are available in the manuscript or the supplementary materials. Correspondence and requests for materials should be addressed to C.-H.H.

## Supplementary Materials

Materials and Methods

Supplementary Text

Figs. S1 to S8

References (*17, 20, 24, 27–34*)

Movie S1

## Supplementary Materials

### Materials and Methods

#### DNA Constructs

The construction of the sensor expression vector has been described (*27*). Constructs were inserted by Gateway LR reactions, into the yeast expression vectors pDR-Flip30, 39, 42-linkers and -GW. The vector of pDR-Flip30 sandwiches the insert between an N-terminal Aphrodite t9 (AFPt9) variant (*28*), with 9 amino acids truncated off the C-terminus, and a C-terminal monomeric Cerulean (mCer) (*3*). pDR-Flip39 sandwiches the inserted polypeptide between an N-terminal enhanced dimer Aphrodite t9 (edAFPt9) and C-terminal fluorescent protein enhanced dimer, with 7 amino acids and 9 amino acids truncated from the N-terminus and C-terminus of eCyan (t7.ed.eCFPt9), respectively. pDRFlip42-linker carries an N-terminal citrine and a C-terminal mCerulean (*29*). The pDR-Flip42-linker vector was digested with *KpnI* (NEB) for insertion of additional linker sequences (Arg-Ser-Arg-Pro-Thr-Arg-Pro-Gly-Glu-Leu-Gly-Thr) to generate pDRFlip42-linker vector. The full-length ORF of NasR, the NIT domain of NasR, or NasR carrying point mutations from *Klebsiella oxytoca* (*17*) in the pDONR221 GATEWAY™ Entry vector were used as sensory domains for creating the nitrate sensors NiMet-NIT, NiMet1.0, NiMet2.0, NiMet3.0, nlsNiMet3.0, or NiMet3.0-NRs. The yeast expression vectors were then created by GATEWAY™ LR reactions between different forms of pDONR221-NasR/NIT and different pDR-FLIP-GWs, following manufacturer’s instructions.

#### Generation of NiMet3.0-NR mutants

The nitrate binding domain of the NasR Entry Clone for NiMet3.0 was altered using the QuikChange Lightning Site-Directed Mutagenesis Kit (Agilent Technologies) according to manufacturer’s instructions to generate the NiMet3.0-NR mutations. Primers for site-directed mutagenesis of NiMet3.0 to create NiMet3.0-NR are as followed: R49A, Forward: 5’-catatgctgcagtgtgcacggggagccagtaat-3’, Reverse: 5’-attactggctccccgtgcacactgcagcatatg-3’; R50A, Forward: 5’-gtacatatgctgcagtgtgaagcgggagccagtaatatctg-3’, Reverse: 5’-cagatattactggctcccgcttcacactgcagcatatgtac-3’; R176A, Forward: 5’-cgcgggtcaggcacgggcgctgg-3’, Reverse: 5’-ccagcgcccgtgcctgacccgcg-3’; R236A, Forward: 5’-gagattgagcagctggctcgtgtcgcttgcac-3’, Reverse: 5’-gtgcaagcgacacgagccagctgctcaatctc-3’.

#### Expression of sensors in yeast

*Saccharomyces cerevisiae* strain BJ5465 [ATCC 208289 (*MATa ura3-52 trp1 leu2-Δ1 his3-Δ200 pep4::HIS3 prb1-Δ1.6 R can1 GAL*] (*30*), obtained from the Yeast Genetic Stock Center (University of California, Berkeley, CA), was transformed with the pDRFlips yeast expression plasmids using a lithium acetate transformation protocol (*31*). Transformed yeast were selected on solid YNB (minimal yeast medium without nitrogen; Difco) supplemented with 2% glucose and - *ura* DropOut medium (Clontech). Single colonies were grown in 5 mL liquid YNB supplemented with 2% glucose and −*ura* drop out under agitation (230 rpm) at 28 °C until OD_600nm_ ~ 0.8 was reached for fluorescence analysis of sensor expression and for metal-affinity chromatography purification of sensors. Yeast strains expressing sensors were grown in 30-ml cultures in −*ura* DropOut medium in 50-ml culture tubes.

#### Fluorescence analysis of purified sensors

Biosensors were purified by metal affinity chromatography. Yeast lysates were diluted 1:2 in 50 mM MOPS, 10 mM imidazole, pH 7.4 and then filtered through a 0.45-μm PES filter and bound to Poly-Prep chromatography columns (Bio-Rad) containing His-Pur Cobalt resin (Bio-Rad). Columns were then washed twice with 50 mM MOPS, 10 mM imidazole, pH 7.4 and eluted in 50 mM MOPS, 150 mM imidazole, pH 7.4. Samples were diluted in 50 mM MOPS, pH 7.4. Fluorescence was measured in a fluorescence plate reader (M1000, TECAN, Austria), in bottom-reading mode using a 7.5 nm bandwidth for both excitation and emission (*32, 33*). Typically, emission spectra were recorded (λ_em_ 470-570 nm, step size, 5 nm). To quantify fluorescence responses of the sensors to substrate addition, 100 μL of substrate (dissolved in 50 mM MOPS buffer, pH 7.4) were added to 100 μL of cells in 96-well flat bottom plates (#655101; Greiner, Monroe, NC). Fluorescence from pDRFlip30 (donor: mCER), 39 (donor: t7.ed.eCFPt9), and 42-linker (donor: mCerulean) was measured by excitation at λ_exc_ 428 nm. Determination of the apparent *K*_d_ of NiMet3.0 for nitrate was performed as described previously (*17*). The purified NiMet3.0 protein was pretreated with 0-20 mM nitrate. Data are reported as mean and s.d. of 3–4 replicates, and each experiment was performed at least three times with similar results. After 15 minutes, buffer was exchanged to 50 mM MOPS, pH 7.4, and fluorescence was analyzed. The emission ratio was subsequently calculated dividing the value of the 530 nm by 488 nm range.

#### Expression of NiMet3.0, NiMet3.0-NR-R176A, and nlsNiMet3.0 in Arabidopsis

The p16 promoter (*24*) from the AT3G60245 gene encoding a 16 S ribosomal subunit was used to drive the nuclear-localized NiMet3.0 fusion biosensor, whereas the CaMV 35S promoter (*34*) was used to drive the NiMet3.0 and NiMet3.0-NR-R176A fusion biosensor in plants. The following construct was inserted into the multiple cloning site of the p16-Kan vector (*20*) : 5’ – a sequence coding for the SV40-derived nuclear localization signal LQPKKKRKVGG (*24*), a sequence coding for Aphrodite; a Gateway cassette including *attR1*, Chloramphenicol resistance gene, *ccdB* terminator gene and *att*R2; a sequence coding for mCerulean (mCer); and a sequence coding for the cMyc epitope tag – 3’, or pZPFlip *UBQ10-KAN* vector under control of the *UBQ10* promoter. The resulting Gateway Destination vectors (p16-FLIPnls30 and pZPFlip30) were then recombined in Gateway LR reactions with NasR or NasR-NR-R176A Entry Clones, resulting in NiMet3.0, NiMet3.0-NR-R176A, and nlsNiMet3.0 expression clones. Transgenic plant lines were generated using the *Agrobacterium* floral dip method as described previously (*28*). Transformants were selected on agar plates containing ½ × Murashige Skoog (MS) medium with Kanamycin.

#### Fluorescence microscopy

*Arabidopsis* seedlings were either grown vertically on ½ × MS agar medium (½ × MS salts without nitrogen, 1% Agar, 0.05% (w/v) sucrose, pH 5.7) plates or germinated on hydroponic medium solidified with 1% agar (Becton Dickinson Biosciences) within cut pipette tips, 5-mm long and 1-mm diameter, that were positioned in an upright position onto a plate with solidified medium for confocal images or light-sheet images, respectively. Plates were stratified for 3 days at 4 °C in the dark prior to being placed in a growth chamber with long-day growth conditions (16 h light/8 h dark cycling, temperature cycling 22 °C day/18 °C night, 67% RH). Seedlings for confocal images were placed in solution containing ½ × MS medium (½ × MS salts without nitrogen, 0.05% sucrose pH 5.7) and prepared for imaging on glass slides. Seedlings for light-sheet microscopy were grown for 3 days in the growth chamber, at which time the root tips had almost reached the lower tip outlet. The tips were plugged into a ~3-cm piece of Fluorinated Ethylene Propylene (FEP) tubing with an inner diameter of 0.115 cm, an outer diameter of 0.195 cm, and wall thickness of 0.04 cm (TEF-CAP, AWG17SW-FEP) and sterilized in 70% ethanol. A closed cultivation system within FEP tubing was used for imaging. Both upper and lower FEP tubes were sealed using gaskets, on which an inlet and outlet tube were inserted into each side of the gaskets and connected to silicon tubing within a pumping perfusion system. To maintain the humidity within the closed cultivation system, the inner sides of the tubing holder have surrounding water reservoirs. Upon transfer to the light-sheet microscope, the seedling was illuminated by a light connected to a timer switch to maintain the light/dark period. The FEP tubing was filled with half-strength hydroponic medium without nitrogen and incubated for 3 days. The FEP tubing was then fixed in a metal holder and placed into the light-sheet microscopic chamber, which was filled with water. The half-strength hydroponic medium without nitrogen (pH 5.5) was continually replaced using a peristaltic pump (GE HealthCare) with a flow rate of 1 mL per hour. The temperature of the microscopy chamber was set at 22 °C.

For nitrate treatments on glass slides, seedlings were placed on glass slides with 50 μL solution and surrounded with a rectangle of vacuum grease and covered with a square cover slip equal in height and half the width of the vacuum grease rectangle. The nitrate treatment solution could then be exchanged beneath the coverslip by addition to the left and removal from the right side of the coverslip. Images were acquired at the time points as indicated in each figure.

Confocal images were acquired on a Zeiss 780 using a 20x/0.8 Plan-Apochromat dry objective or 40x/1.2 C-Apochromat water objective. Cyan fluorescent protein (CFP; 440 nm) and YFP (514 nm) were excited with lasers. Fluorescence emission was detected by a GaAsP PMT detector, set to detect 463 to 508 nm for CFP, and a normal PMT detector, set to 520 to 585 nm for YFP. The laser power was set between 0.5% and 2% with detector gain set to 700-750 to image CFP or YFP.

The lab-established light-sheet system coordinated with a *micro*LAMBDA Pte Ltd (Singapore). Light-sheet imaging was performed using a 20x 0.5 dipping objective; two illumination arms with galvanometer scanners; 10x long working distance objectives; and 445 nm and 515 nm lasers used for excitation of CFP and YFP, respectively. For FRET measurements, sequential imaging of cyan and yellow fluorescent proteins is performed with a DC filter wheel with ET470/24m and an ET535/50m emission filters, driven by a MAC6000 controller (Ludl Electronic Products, Hawthorne, NY). Fluorescence emission was detected by a Hamamatsu Flash 4.0 V3 camera. Imaging data were acquired using MetaMorph software (Downingtown, PA). Data were taken as time series with simultaneous acquisition of FRET donor and acceptor fluorophores under donor excitation, followed by acquisition of donor and acceptor under acceptor excitation.

#### Image processing and analysis

Image processing and fluorescence pixel intensity were quantified using Fiji software (http://fiji.sc/). Mean gray values of regions of interest within the root meristem region were calculated as follows: Background was subtracted from all measured intensities as generated ROIs where there was no plant material, measured mean intensity values in all four channels (Dx/Dm; Dx/Am; Ax/Dm; Ax/Am) and subtracted that intensity from the entire image. Ratio images (DxAm/DxDm) were created using the Ratio Plus plug-in for ImageJ (Paulo Magalhães, University of Padua, Italy). Regions of interest were selected and analyzed with the help of the ROI manager tool.

In this work, we presented data using Beeswarm and box plots of raw data. In the beeswarm and box plot graphs, the central rectangle spans the first quartile to the third quartile, while the line inside the rectangle shows the median. The whiskers denote 1.5 interquartile ranges from the box and outlying values plotted beyond the whiskers. All the statistical analyses were performed using GraphPad Prism version 9.0.0 for MAC (www.graphpad.com).

## Supplementary Text

**Fig. S1.**
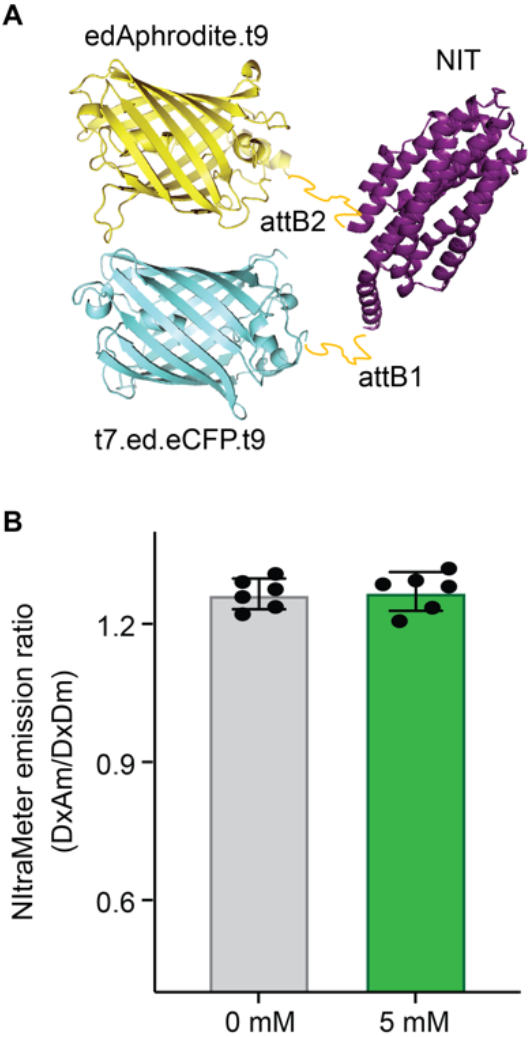
Fluorescence emission ratio of NiMet-NIT. **A**, Structural model of NiMet-NIT. NIT was fused via *attB1* and *attB2* linkers to fluorescent protein FRET pairs (donor, edAphrodite.t9, and acceptor, t7.ed.eCFP.t9, fluorescent proteins). The NIT protein (purple) representation is from a published structure [PDB 4AKK(*17*)]. **B**, Purified NiMet-NIT proteins with and without 5 mM NO_3_^-^ treatment. Mean and s.d. of six biological repeats are presented.

**Fig. S2.**
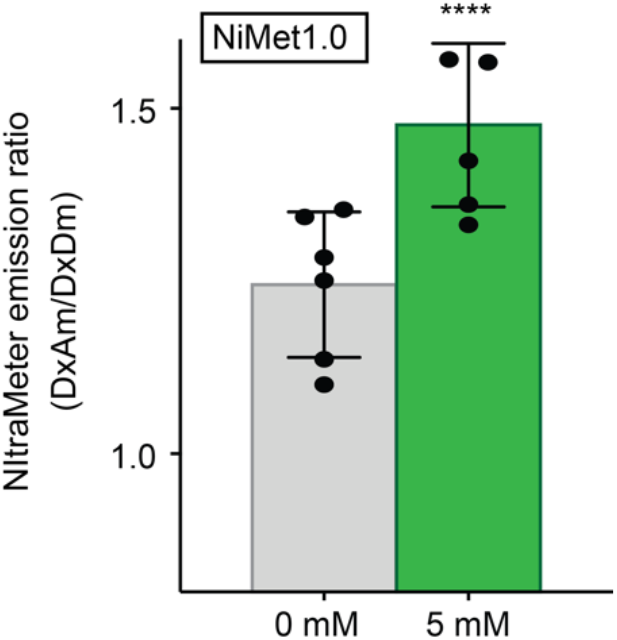
Fluorescence emission ratio of NiMet1.0. NasR was fused via *attB1* and *attB2* linkers to fluorescent protein FRET pairs (donor, edAphrodite.t9, and acceptor, t7.ed.eCFP.t9, fluorescent proteins). Purified NiMet1.0 proteins with and without 5 mM NO_3_^-^ treatment. Student’s *t*-test \*\*\*\**P* value < 0.0001. Mean and s.d. of six biological repeats are presented.

**Fig. S3.**
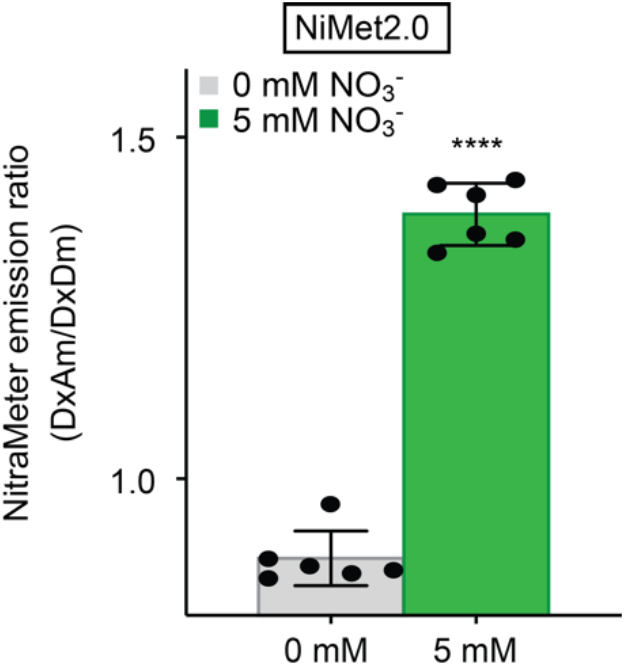
Fluorescence emission ratio of NiMet2.0. NasR was fused via *attB, attB2*, and L12 linkers to fluorescent protein FRET pairs (donor, Citrine, and acceptor, mCerulean, fluorescent proteins). Purified NiMet2.0 proteins with and without 5 mM NO_3_^-^ treatment. Student’s *t*-test \*\*\*\**P* value < 0.0001. Mean and s.d. of six biological repeats are presented.

**Fig. S4.**
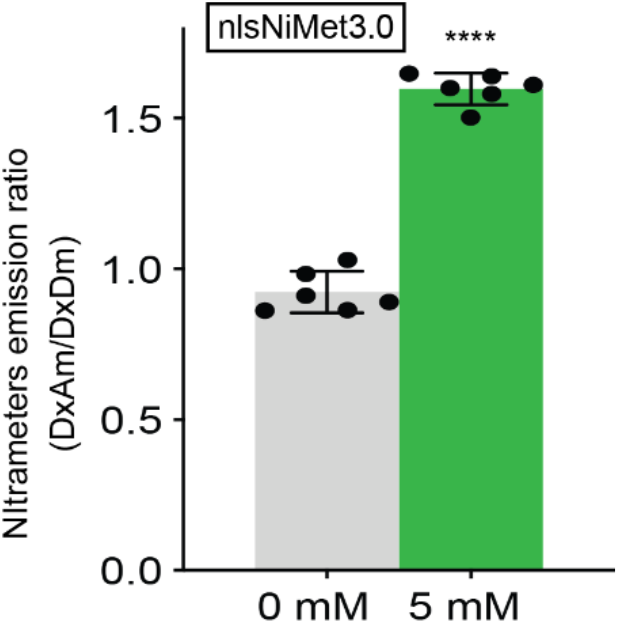
Fluorescence emission ratio of nlsNiMet3.0. Purified nlsNiMet3.0 proteins with and without 5 mM NO_3_^-^ treatment. Student’s *t*-test \*\*\*\**P* value < 0.0001. Mean and s.d. of six biological repeats are presented.

**Fig. S5.**
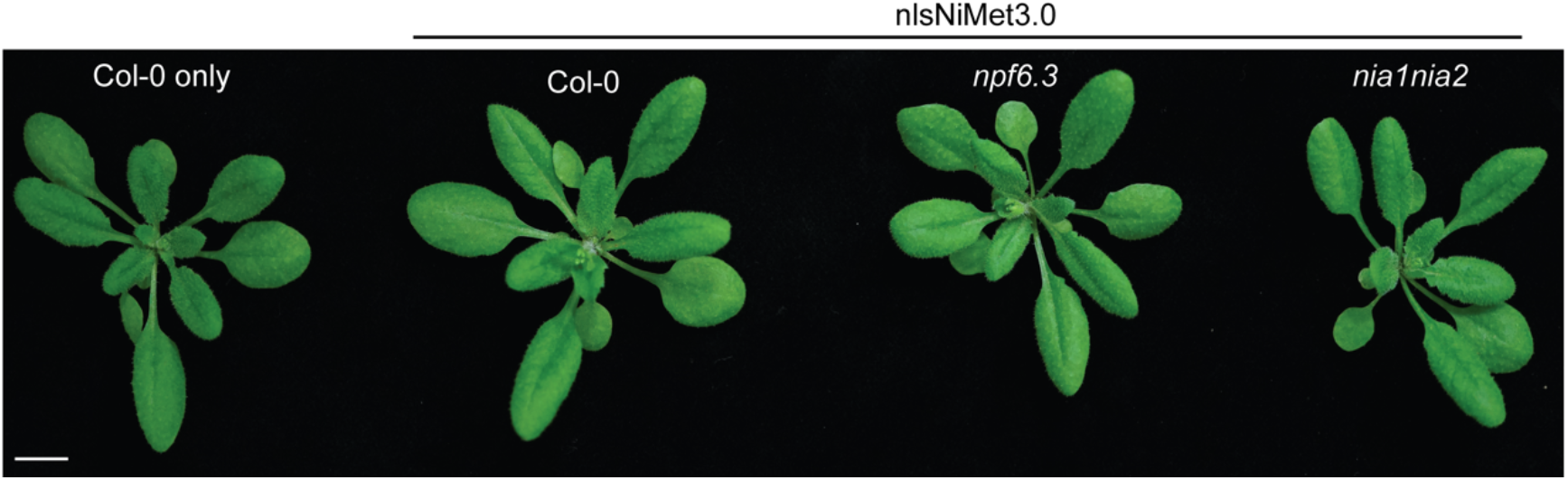
Seedlings of Col-0 and mutants expressing nlsNiMet3.0. No significant growth phenotype was observed in plants expressing nlsNiMet3.0 in Col-0, *npf6.3* or *nia1nia2* backgrounds plants when compare with wild type Col-0. Pictures were taken while plants were grown under full nutrient conditions, day/night cycles 16/8 hours for 14 days. Scale bar, 1 cm.

**Fig. S6.**
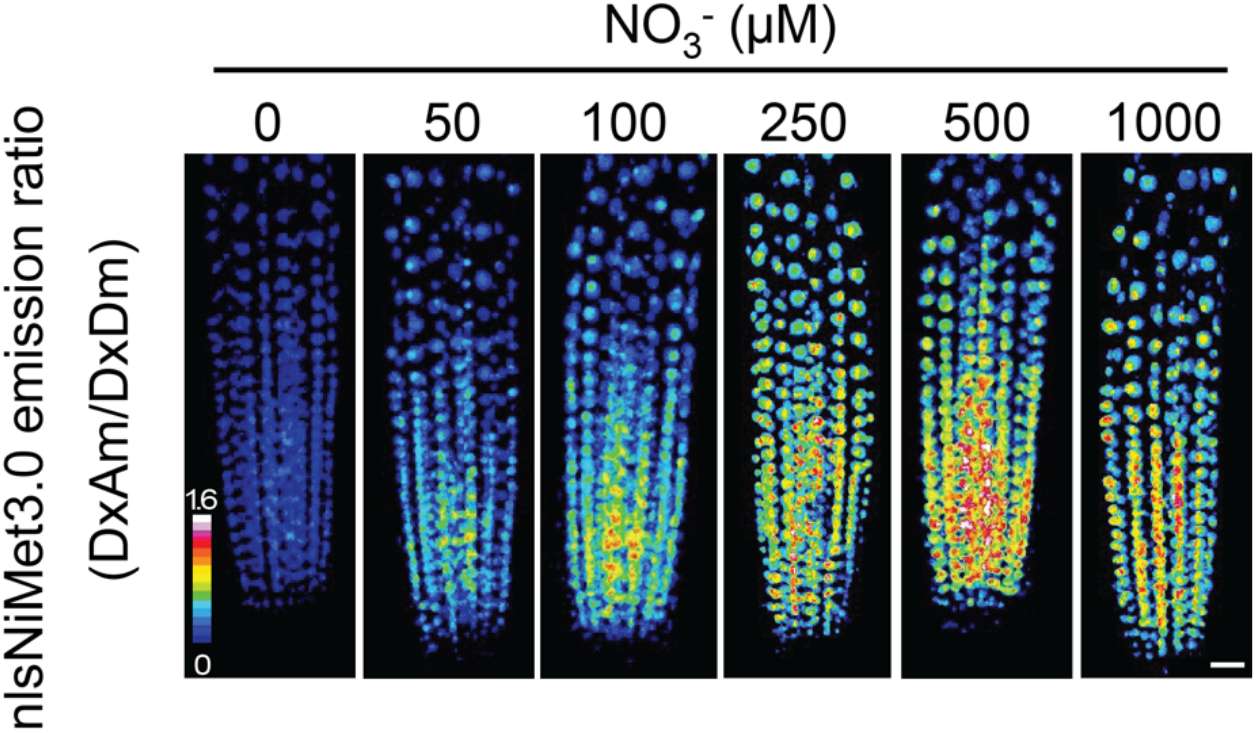
S6 nlsNiMet3.0 fluorescence response to increasing concentrations of NO_3_^-^ in root. Three-dimensional images of nlsNiMet3.0 emission ratios in 6-day-old roots in Col-0 background. The concentrations of NO_3_^-^ are indicated in figure. Scale bar, 25 μm. Complete experiments were repeated at least three times with similar results.

**Fig. S7.**
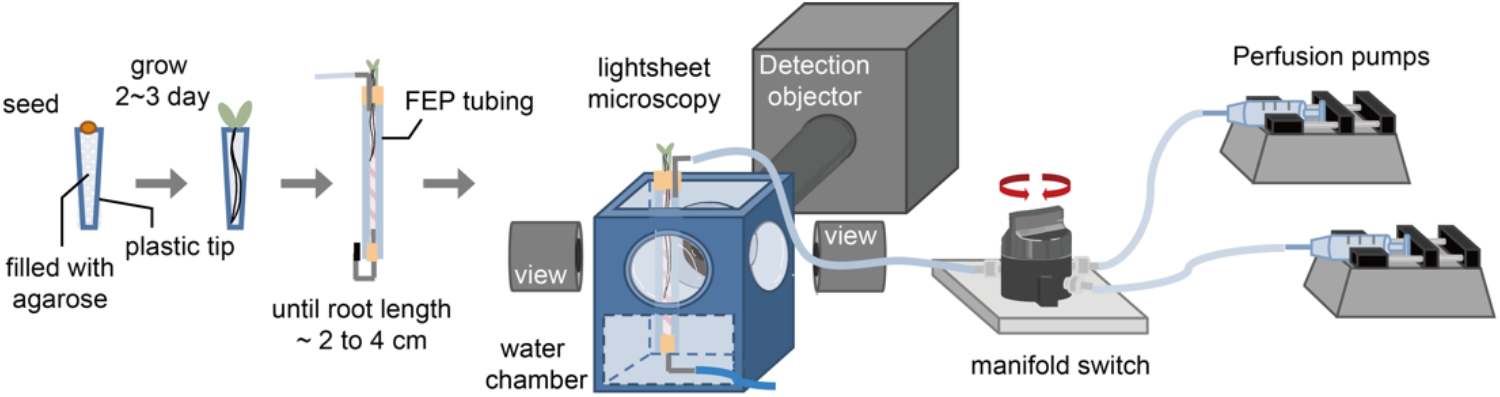
Workflow for imaging Arabidopsis root in light-sheet microscope with perfusion system. Sample workflow for *in vivo* microscopic imaging in lab-designed light-sheet apparatus. *Arabidopsis* seeds are germinated on top of nutrient agarose solidified in a plastic tip. Plastic tips are placed on one end of a piece of FEP tubing, where the root tip is subjected to liquid flow and becomes accessible for imaging. An optional valving system provides precise control over the flow during experiments where conditions need to be changed rapidly. Imaging then occurs on a light-sheet microscope.

**Fig. S8.**
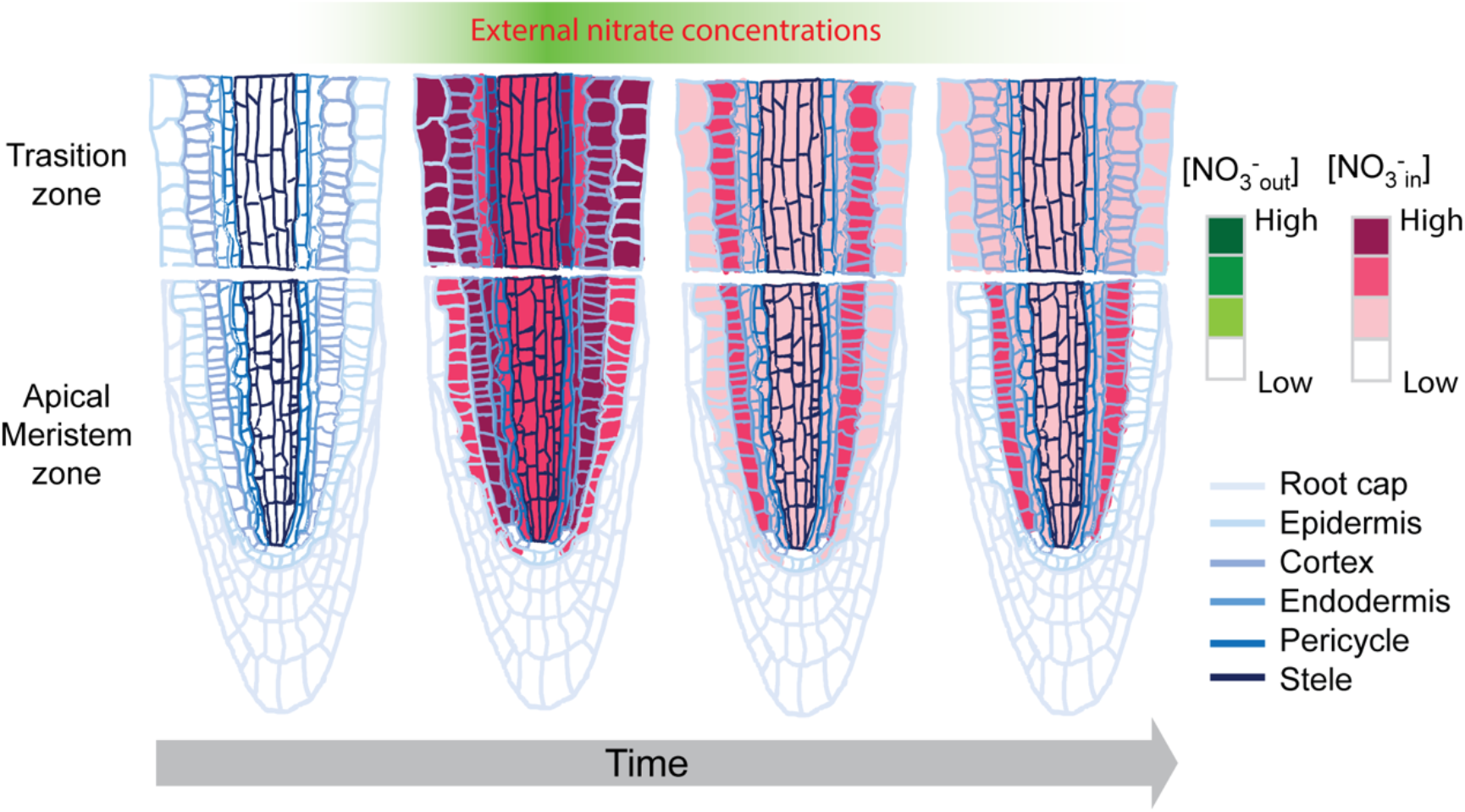
Nitrate distribution in the root before and after treatment with exogenous NO_3_^-^. The NO_3_^-^ distribution in the meristem and transition zones is based on the results in the Fig. 4. [NO_3_^-^], indicates the concentration of NO_3_^-^. in, indicates inside the cells; out, indicates outside the cells.

**Movie S1.**

The movie shows a time-course treatment of nlsNiMet3.0 in root with NO_3_^-^. The movie shows the emission ratio (DxAm/DxDm). Scale bar, 25 μm.

